# Anthropogenic noise affects *Oikopleura dioica* behaviour in a life stage- and signal-dependent manner

**DOI:** 10.64898/2026.11.19.767843

**Authors:** Eva R. Quintana, Oleg Tolstenkov, Rodolfo da Silva Mazzarini Baldinotti, Sissel Norland, Valentina Zaffaroni-Caorsi, Ignasi Nou-Plana, Lucia Manni, Giacomo Sabbadin, Simona Candiani, Matteo Bozzo, Emanuele Serafini, Roberta Pennati, Giorgio Blumer, Irene Guarneri, Filomena Ristoratore, Anne Elin Aasjord, Daniel Marc Chourrout, Rosa Maria Alsina-Pagès, Giovanni Zambon, Cristian Cañestro, Marios Chatzigeorgiou

## Abstract

Anthropogenic noise is an increasingly important pollutant of marine environments, yet its effects on zooplankton remain poorly understood. The appendicularian *Oikopleura dioica* is a globally distributed filter-feeding tunicate whose locomotion and house-building behaviour directly influence feeding efficiency and carbon export. Here, we investigated how anthropogenic noise affects behaviour across the life cycle of *O. dioica*, from larvae to juveniles and adults, using stage-specific tracking and quantitative behavioural analyses to assess responses to a high-intensity standardized noise band (MSFD-NB) and a vessel-derived noise signal (Vessel-DNS). Animals exhibited consistent behavioural alterations during and after noise exposure, and the responses differed strongly among life stages and between noise signals. Larvae showed the greatest sensitivity, displaying increased locomotor activity and altered movement dynamics during exposure, as well as persistent post-exposure effects. Juveniles exhibited treatment-dependent changes in trajectory organization, whereas adults showed modifications in orientation, turning behaviour, and tail-driven propulsion. In adults residing within their filter-feeding house, noise altered pumping-associated movements without affecting house-expansion frequency.

These findings demonstrate that anthropogenic noise impacts behaviour throughout the *O. dioica* life cycle in a stage- and signal-specific manner, suggesting that noise pollution may not only impact individual performance, but potentially also ecological processes linked to marine food webs, carbon sequestration, and consequently, ocean health.

## INTRODUCTION

Anthropogenic noise has become one of the most pervasive and fastest-growing forms of pollution in the ocean. The expansion of human activities at sea has driven a substantial and sustained rise in underwater noise levels, transforming marine soundscapes at an unprecedented rate^1–3^. These effects are documented across multiple levels of biological organisation, from molecular and physiological stress responses^4^ to changes in behaviour^5^ and, ultimately, population-level consequences^6^. In this context, behavioural responses are among the most frequently documented consequences and often constitute the earliest observable indicators of disturbance. These responses range from subtle disruptions in communication^7^ and avoidance responses^8^ to overt effects on feeding, reproduction, and social interactions^7,9,10^. Because affected species often occupy key ecological roles, ranging from primary consumers to top predators, understanding how noise reshapes their behaviour is essential for anticipating broader ecosystem consequences.

Although considerable progress has been made in understanding the effects of anthropogenic noise on marine organisms, substantial gaps remain, particularly for marine invertebrates. Zooplankton are ecologically important because they occupy a central position in marine food webs and biogeochemical cycles, mediating energy transfer from primary producers to higher trophic levels and contributing substantially to carbon cycling and ecosystem stability^11,12^. Noise-induced changes in their survival, locomotion or filter feeding could therefore generate cascading bottom-up effects. In copepods, for instance, acoustic stress can increase mortality, delay development and reduce body size^6,13,14^. These effects are not universal: comparable airgun exposure produced no detectable changes in mortality, behaviour or gene expression in *Calanus finmarchicus*^15^, indicating that susceptibility depends on species, life stage and exposure regime. However, locomotor and feeding behaviours, which directly govern prey capture, predator avoidance and ultimately survival and reproduction in small pelagic invertebrates, remain comparatively understudied.

Among filter-feeding zooplankton, tunicates clearly illustrate this gap, in which noise-induced behavioural effects in this group remain almost entirely uncharacterised. Existing work has largely focused on the acoustic sensitivity of larval settlement and metamorphosis cues of ascidians^16^; direct behavioural responses to acoustic treatment in adults have only recently begun to be characterised, with siphon-withdrawal responses to noise in *Styela plicata* and ultrasound reported across several solitary ascidian species^17,18^ and substrate-borne vibration, rather than sound pressure, identified as the effective stimulus in the ascidian *Halocynthia papillosa*^19^.

Appendicularians such as *Oikopleura dioica* provide a powerful framework for investigating how environmental stressors influence behavioural organization throughout development. In addition to their central role in pelagic food webs through the larvacean shunt^20,21^, these animals contribute disproportionately to carbon cycling through the continuous production and discard of their filter-feeding houses^22^, which aggregate into marine snow and drive a substantial share of vertical carbon export^12,23^. Because both ecological functions depend directly on sustained feeding and locomotor activity, noise-induced disruption of behaviour in *O. dioica* has the potential to affect ecosystem-level processes well beyond the individual.

*O. dioica* is a small, fully pelagic tunicate that completes its entire life cycle, from fertilisation to sexual maturity, in approximately seven days under the conditions used here (12 ± 0.5 °C)^24^. Unlike other tunicates, *O. dioica* remains free-living throughout its life and continuously builds and discards a transparent gelatinous “house” that it uses to filter-feed from the surrounding water column. *O. dioica* possesses specialised mechanosensory structures, including the circumoral ring (also termed the coronal organ) composed of ciliated sensory cells in the mouth associated with feeding-related particle rejection, and the posteriorly located Langerhans receptors implicated in mechanically evoked escape responses^25,26^. These structures provide a basis for investigating the animal’s potential sensitivity to mechanical or acoustic disturbance, although their contribution to responses to environmental noise remains unclear.

By examining noise-induced changes in the stereotypical behavioural repertoire of *O. dioica*, we aimed to address the limited understanding of how acoustic pollution affects zooplanktonic organisms and whether these responses may translate into functional consequences for feeding and locomotion, which ultimately may affect animal survival and reproduction.

Previous work has shown that behavioural maturation in this species is not driven by the emergence of novel motor behaviours, but by progressive changes in the coordination, kinematics, posture usage and transition dynamics of a conserved behavioural repertoire^27^. Larvae show relatively simple, episodic swimming, while juveniles exhibit more sustained locomotion. Adults display the most complex trajectories, further shaped by house-associated constraints on movement^27^.

Building on this framework, we tested under controlled laboratory conditions how *O. dioica* responds behaviourally to two distinct anthropogenic noise signals: a high-intensity standardized noise band (MSFD-NB) and a vessel-derived noise signal (Vessel-DNS). Specifically, we asked whether (i) anthropogenic noise exposure alters the stereotyped behavioural repertoire of *O. dioica*; (ii) these effects differ in magnitude or nature among larvae, juveniles, and adults as sensory structures and locomotor capacities mature; and (iii) either noise signal disrupts house-associated behaviour, given the house’s direct and continuous exposure to the hydrodynamic environment and its central role in filter-feeding.

## RESULTS

To study the impact of anthropogenic noise on behavioural development in the appendicularian *O. dioica*, we recorded videos of larvae, juveniles and adults (inside or outside the house) using previously described tracking platforms^27–29^.

Behaviour was assessed under two complementary acoustic treatments (see Methods, Table S1 and S2). High-intensity experimental noise, following EC-MSFD guidelines (MSFD-NB, Marine Strategy Framework-Noise Band^30^), was first used to characterise the potential effects of strong sustained acoustic treatment. Vessel-signal noise representative of maritime traffic in European basins (Vessel-DNS, Vessel-Deuteronoise Sound^31^) was subsequently used to determine whether similar responses could occur under an environmentally grounded vessel-derived signal.

### Behavioural responses during noise exposure across ontogeny

To determine whether *O. dioica* altered its swimming behaviour during acute exposure, we analysed larvae (7.5 h post-fertilisation (hpf) at 14 °C), juveniles (18 hpf at 12 °C) and adults (days 4-5 of the 7-day life cycle). Separate cohorts were used for the MSFD-NB and Vessel-DNS experiments; animals were not reused across treatments. Larvae and juveniles were recorded for 15 min before, during and after a 15-min exposure period. The exposure segment was analysed and compared with separately recorded controls. Adults were not recorded before exposure because they could begin expanding a new house after removal from the original house. We quantified animal-level kinematic, trajectory and postural metrics. Representative metrics were selected through feature filtering and principal component analysis (PCA) to identify informative, non-redundant features capturing the main dimensions of behavioural variation (Fig. S1; Tables S10-S11) and compared between each noise-exposed group and its corresponding, separately recorded control group (Fig. 1; Tables S3-S5).

**Figure 1.**
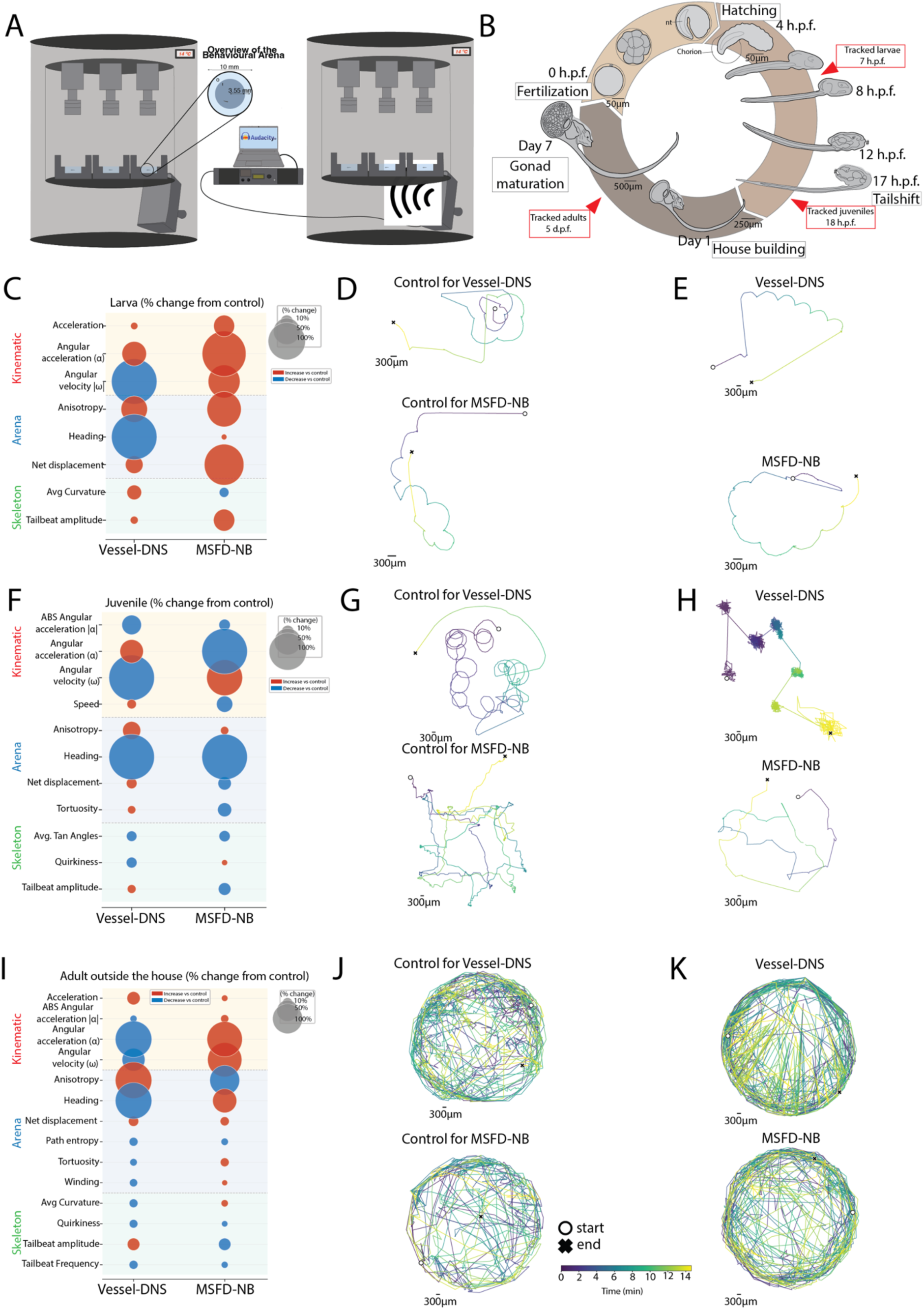
Behavioural responses of larvae, juveniles and adults (outside the house) during MSFD-NB and Vessel-DNS exposure, relative to matched controls. **(A)** Overview of behavioural recording set-up as detailed in [28, 29], including the noise-exposure set-up and overhead view of the arena. Further details are provided in the Materials and Methods. (B) *O. dioica* life cycle at 12–14 °C and tracking times for each life stage. (C, F, I) Percentage change from control in kinematic, arena and skeleton parameters for larvae, juveniles and adults, respectively; circle size indicates magnitude and colour indicates direction (red, increase; blue, decrease). (D-E, G-H, J-K) Representative trajectories for control, MSFD-NB and Vessel-DNS groups in larvae, juveniles and adults, respectively; colours indicate elapsed time. Parameter definitions and sample sizes are provided in Supplementary Tables S1 and S2, respectively. Metrics were averaged per animal before analysis. Statistical significance was assessed using two-sided Mann–Whitney U tests with Benjamini–Hochberg (BH) correction across metrics within each stage and noise treatment.. Unadjusted and BH-adjusted p-values and effect sizes are provided in Tables S3-S5. Scale bars, 300 µm.

### Acute behavioural responses in larvae during noise exposure

During MSFD-NB exposure, larvae showed significant differences in several behavioural metrics relative to larvae in the corresponding, separately recorded control group (Fig. 1C; Table S3, Fig. S3 for features statistical overview). Their overall multivariate behavioural profile also differed significantly from that of the control group (Fig. S2A; Table S12). MSFD-NB-exposed larvae showed increases in net displacement, swimming speed, anisotropy and tail-beat amplitude, together with lower winding, angular velocity and angular acceleration (Fig. 1C; Fig. S3; Table S13). Collectively, these changes indicate that the exposed larvae moved faster and more directionally, covering a greater net distance along less convoluted trajectories than control larvae. This combination of increased anisotropy and reduced winding may reflect reduced behavioural variability and more stereotyped swimming, although the present analysis cannot determine whether it represents a stress-induced escape response. Consistent with these quantitative measurements, the representative trajectory plots show greater net displacement and less recurrent exploration of the arena by MSFD-NB-exposed larvae relative to control larvae (Fig. 1D–E).

Vessel-DNS-exposed larvae showed less pronounced differences from their corresponding control group. They exhibited tendencies towards greater anisotropy and net displacement, together with reduced changes in heading and turning dynamics (Fig. 1C; Table S3). These patterns were consistent with more directional swimming and altered reorientation within the arena relative to control larvae. However, none of these differences remained significant after correction for multiple comparisons (Figs. S2–S3; Tables S12–S13). Therefore, although the descriptive patterns observed under Vessel-DNS differed from those observed under MSFD-NB, no direct statistical comparison between the two acoustic treatments was performed.

### Signal-specific changes in juvenile trajectories

Juveniles exposed to either acoustic treatment showed no behavioural differences from their corresponding control groups that remained significant after correction for multiple comparisons, in contrast to the significant differences observed in MSFD-NB-exposed larvae. During MSFD-NB exposure, juveniles showed tendencies towards lower tortuosity, altered angular acceleration and velocity, and greater anisotropy relative to their corresponding controls (Fig. 1F–H; Table S4). Collectively, these patterns were consistent with more directional and less irregular swimming trajectories. However, none of the differences remained significant after correction for multiple comparisons (Fig. S3; Table S13).

During Vessel-DNS exposure, juveniles showed tendencies towards altered turning dynamics, reduced heading and increased tortuosity relative to their corresponding controls (Fig. 1F; Table S4). These patterns were consistent with more sinuous trajectories and altered reorientation during Vessel-DNS exposure (Fig. 1G-H), rather than a general increase or decrease in locomotor activity. Again, none of these differences remained significant after correction (Figs. S2–S3; Tables S12–S13). The juvenile results should therefore be interpreted as treatment-associated tendencies rather than robust behavioural effects. No direct statistical comparison was performed between the MSFD-NB and Vessel-DNS treatments.

### Changes in free-swimming adult behaviour

Free-swimming adults outside the house showed treatment-associated differences across several behavioural features relative to their corresponding control groups, although none remained significant after correction for multiple comparisons (Figs. S2–S3; Tables S12–S13). During MSFD-NB exposure, adults showed tendencies towards increased heading and angular acceleration, reduced anisotropy, and lower tail-beat amplitude and frequency relative to their corresponding controls (Fig. 1I; Table S5). These patterns may indicate changes in trajectory organisation and tail-driven propulsion, but the representative trajectories did not show a sufficiently clear difference between exposed and control adults to support further interpretation (Fig. 1J–K).

During Vessel-DNS exposure, adults showed tendencies towards increased acceleration and anisotropy, decreased heading and angular velocity, higher tail-beat amplitude, and lower tail-beat frequency relative to their corresponding controls (Fig. 1I; Table S5). Collectively, these patterns may reflect changes in the directional, rotational and propulsive components of swimming. However, none remained significant after correction for multiple comparisons. Adult responses varied substantially among individuals and should therefore be considered descriptive, treatment-associated tendencies rather than robust behavioural effects.

Overall, behavioural responses during exposure were life-stage dependent. Most differences that remained significant after correction for multiple comparisons were observed in larvae exposed to MSFD-NB, whereas juveniles and free-swimming adults showed only non-significant tendencies in trajectory organisation and propulsion. Vessel-DNS exposure produced no significant differences from the corresponding controls at any life stage. Under the conditions tested, these findings suggest that larvae are more sensitive, and potentially more vulnerable, to acute acoustic exposure than older life stages. This greater sensitivity may reflect a vulnerable period of development or a more limited capacity to compensate for external stressors. Conversely, the absence of significant effects in juveniles and adults may reflect greater resilience or compensatory capacity, potentially associated with the continued development of sensory and motor systems. These interpretations remain hypotheses and require direct experimental testing.

Thus, behavioural responses to anthropogenic noise in *O. dioica* appear to depend on both life stage and the acoustic characteristics of the playback signal.

### Changes in house-associated adult behaviour

A previous study has shown that, under normal conditions, in-house adults (Fig. 2A) naturally exhibit distinct behavioural patterns compared to free swimming adult individuals^27^. Therefore, we wanted to investigate which in-house behavioural changes occurred during acute noise exposure compared with the in-house control condition.

**Figure 2.**
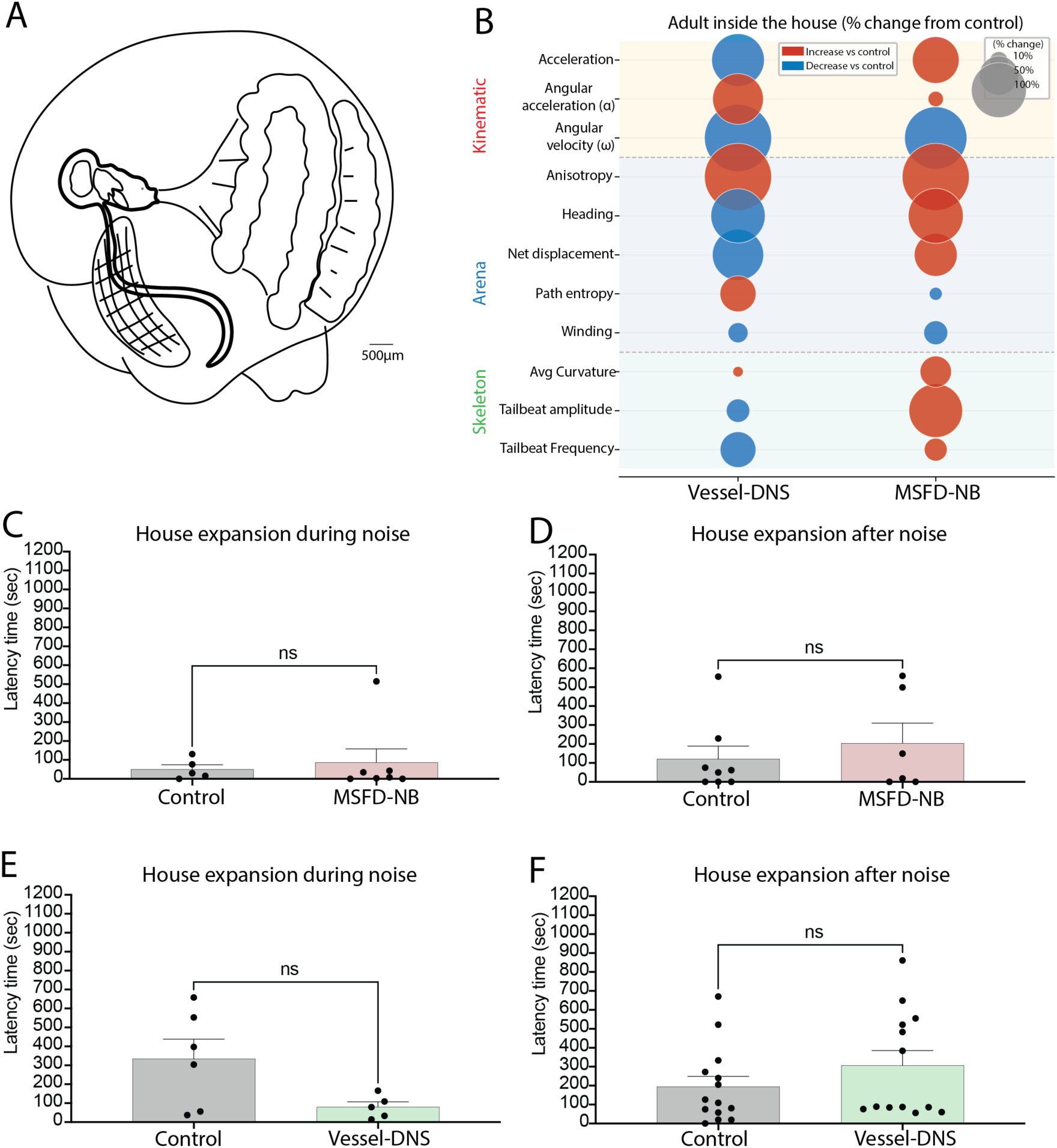
Adult in-house behaviour and house-expansion latency following MSFD-NB and Vessel-DNS exposure. **(A)** *O. dioica* trunk and house, illustrating the animal’s position within its filtering structure. **(B)** Percentage change from control for kinematic, arena and skeleton parameters of adults inside the house. Circle size indicates the magnitude of change and colour indicates direction (red, increase; blue, decrease). **(C, D)** House-expansion latency (s) during and after MSFD-NB noise, respectively, relative to control. **(E, F)** House-expansion latency (s) during and after Vessel-DNS noise, respectively, relative to control. Parameter definitions and sample sizes are provided in Supplementary Tables S1 and S2, respectively. Metrics were averaged per animal before analysis. Noise-versus-control comparisons used two-tailed Mann–Whitney U tests. Behavioural metric p-values were BH-adjusted within each stage, assay, and noise treatment. Unadjusted and BH-adjusted p-values are reported in Table S6; “ns” indicates not significant.

During MSFD-NB treatment, adults inside the house tended to show higher acceleration, stronger body curvature, increased tail-beat amplitude, and changes in heading and net displacement compared to control (Fig. 2B). Within the house, body bending and tail-beats are thought to be directly linked to water pumping and filtration. Therefore, the combined increase in curvature and tail-beat amplitude suggested that MSFD-NB noise could alter the intensity or pattern of pumping-related movements. This differed from adults outside the house, where tail-beat related variables tended to decrease, indicating that noise may affect free swimming and house-associated movement in different ways. This pattern showed a different tendency under Vessel-DNS treatment (Fig. 2B; Table S6), where exposed individuals showed significantly reduced net displacement, together with tendencies towards lower acceleration, lower tail-beat amplitude and frequency, and changes in angular velocity. Together, these changes were consistent with reduced body and tail movement under these noise conditions.

Latency to house expansion did not differ significantly from controls during or after exposure under either noise treatment (Fig. 2C–F). Thus, while movement patterns inside the house showed treatment-associated tendencies, there was no clear evidence of altered house-expansion behaviour.

### Behavioural responses after noise exposure across ontogeny

To determine if noise-driven behavioural changes in *O. dioica* persisted after prolonged noise exposure larval, juvenile and adult (*outside the house*) trajectories were recorded for 15 min after prolonged noise exposure (2 h for larvae, 1 h for juveniles and adults). Even after long exposure, analysis revealed stage-specific behavioural changes observed in kinematic, trajectory and postural metrics.

Larvae again showed the most pronounced post-exposure response. Following prior MSFD-NB exposure, larvae showed higher tail-beat amplitude and angular velocity, combined with a decrease in acceleration, indicating stronger tail beating and faster turning in swimming once the noise had stopped (Fig. 3B; Table S7). Tail-beat amplitude and absolute angular velocity remained significant after BH correction, whereas the decrease in acceleration did not.

**Figure 3.**
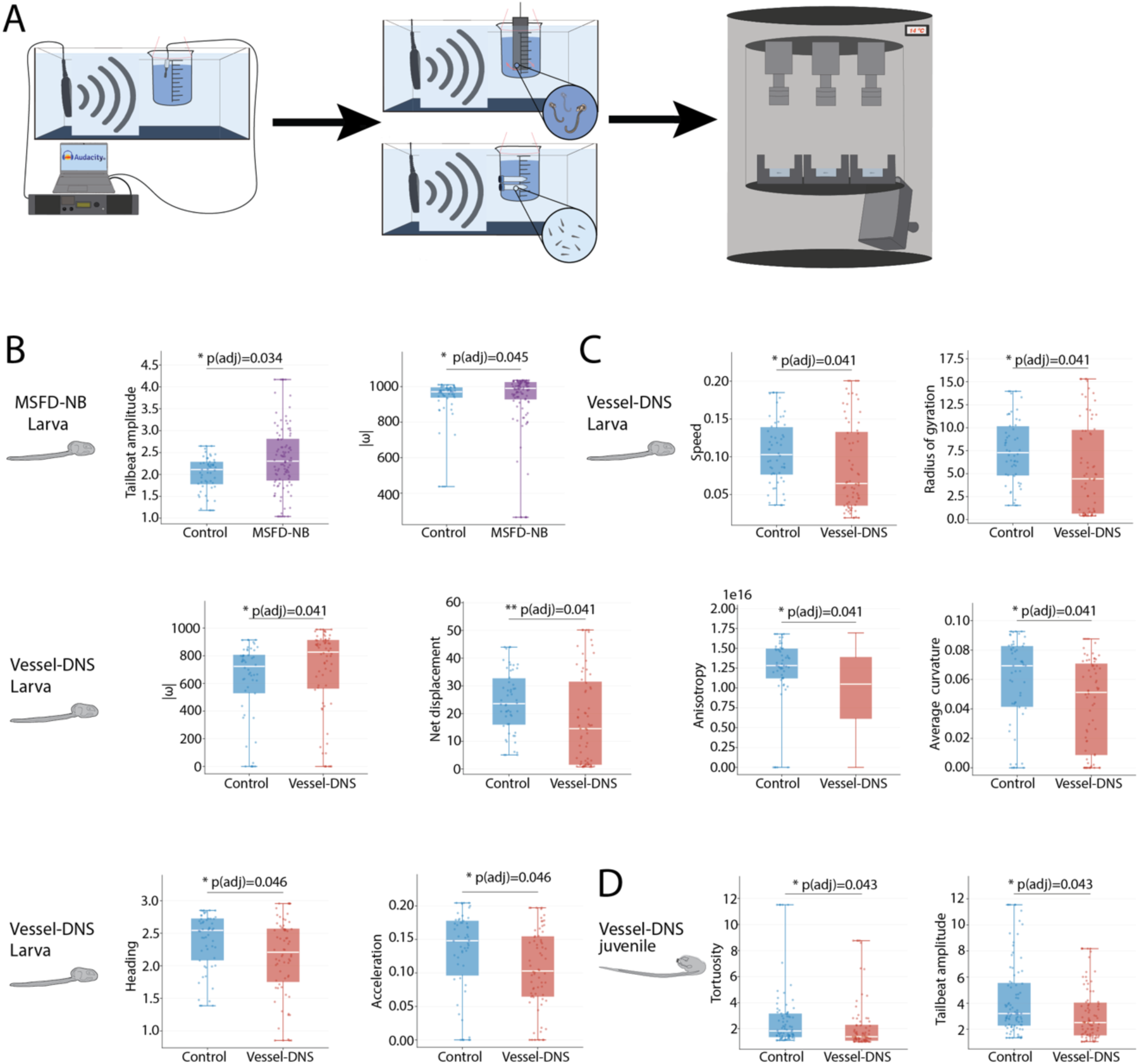
Behavioural responses of *O. dioica* larvae and juveniles after MSFD-NB and Vessel-DNS exposure, relative to matched controls. Box plots show selected behavioural parameters showing differences between noise-exposed animals and their respective controls. **(A)** Overview of behavioural recording set-up as detailed in [28, 29]; further details are provided in the Materials and Methods. **(B)** MSFD-NB larvae: tail-beat amplitude and |ω| quantification **(C)** Vessel-DNS larvae: speed, radius of gyration, |ω|, net displacement, heading, average curvature, anisotropy and acceleration parameters quantification in control and vessel-DNS pre-exposed larvae. **(D)** Vessel-DNS juveniles: tortuosity, tail-beat amplitude quantification in control and Vessel-DNS juveniles. Parameter definitions and sample sizes are provided in Supplementary Tables S1 and S2, respectively. Points represent individual animals; boxes show the median and IQR, and whiskers extend to the 5th and 95th percentiles. Noise-versus-control comparisons used two-tailed Mann–Whitney U tests; BH correction was applied across all 19 behavioural metrics within each developmental stage and noise treatment. Both unadjusted and BH-adjusted p-values are reported in Tables S7-S9(*p < 0.05; **p < 0.01).

After Vessel-DNS exposure larvae showed decreased swimming speed, acceleration, net displacement and radius of gyration, indicating an overall reduction in locomotor output. Their paths were also less directional, less elongated, with reduced anisotropy, lower body curvature and altered heading (Fig. 3C; Table S8). Together, these changes suggested that larvae travelled shorter, slower and less directional paths after the treatment stopped while showing altered turning dynamics.

Juveniles showed a limited change after MSFD-NB exposure. In contrast to larvae, they showed lower tail-beat amplitude (Table S13), a tendency towards reduced tail movement, although this difference was not significant after correction. As in larvae, juveniles showed more differences from controls after Vessel-DNS than after MSFD-NB exposure: juveniles showed reduced tail-beat amplitude and reduced path tortuosity, both significant after correction, a shift in heading and increased anisotropy that were significant only before correction (Fig. 3D; Table S9), indicating reduced tail movement and straighter swimming trajectories after Vessel-DNS exposure. Adults showed no persistent significant changes after noise exposure under any treatment (Table S13).

Together, these results suggested that noise can leave at least a short-term behavioural trace in *O. dioica* developmental stages after the signal has stopped. Importantly, this response was both stage and signal specific and was not simply a continuation of the acute response during the treatment. Juveniles retained a more limited change in propulsion, consistent with a carry-over effect on movement organisation while larvae showed the clearest sensitivity, which potentially reflected the relevance of particle-motion cues for small-bodied early stages.

## DISCUSSION

Understanding how environmental stressors, including the increasing prevalence of underwater noise pollution, affect marine invertebrates is essential for assessing the vulnerability of marine ecosystems and, ultimately, ocean health. Zooplankton are particularly informative in this regard, given their considerable influence on energy transfer within pelagic food webs and their contribution to the biological carbon pump, to which appendicularians contribute a disproportionate share of the vertical carbon flux, up to 55% in coastal systems and up to 39% of total export in mesocosm experiments with *O. dioica* itself^12,23,32^.

Appendicularians contribute to carbon export through their filter-feeding houses, which capture particles and, when discarded, contribute to sinking marine snow^22,23^. Noise-induced changes in feeding and house-associated pumping can therefore affect behaviours directly linked to carbon export, although carbon export was not measured in this study. Within this group, appendicularian tunicates such as the cosmopolitan *O. dioica* provide an attractive model for studying these perturbations because of their rapid life cycle, tractability in laboratory culture and stereotyped behaviour through ontogeny^27,33,34^.

Previous work demonstrated morphological resilience to noise exposure across *O. dioica* development^35^, despite pronounced transcriptomic responses during embryogenesis. Because zooplankton such as *O. dioica* rely on precise locomotor control for feeding, predator avoidance and reproductive success, understanding how noise alters behaviour is critical for interpreting vulnerability to this form of pollution. To our knowledge, this is the first study to investigate the effects of noise exposure on *O. dioica* behaviour, connecting known molecular and functional responses to their locomotor basis.

### *O. dioica* behavioural responses to noise across life stages and playback signals

Noise exposure did not simply increase or decrease activity in *O. dioica* larvae; instead, it altered movement structure, revealing a sensitive sensorimotor window during early free-swimming development. Noise-related disruption of larval swimming and orientation has also been reported in other planktonic taxa, including coral planulae, in which boat noise abolishes soundscape-based habitat selection and may affect recruitment^36^. Particle motion is an important component of underwater acoustic fields for small-bodied aquatic organisms^37^: in copepods, the first antennae respond to water displacements of the order of nanometres^38,39^, and in molluscs and crustaceans sound detection depends on particle motion and substrate-borne vibration rather than on pressure^40–42^. *O. dioica* larvae may therefore be affected by the local water motion generated by the acoustic field, although the sensory basis of this response remains unclear. At the larval stage examined (7.5 h post-fertilisation at 14 °C), the circumoral ring and Langerhans ciliated cells are not expected to have reached their mature morphology. These structures become morphologically distinct from the late hatchling stage and are generally considered functional from the tail shift stage onwards (∼19 h post-fertilisation at 13 °C), when juveniles begin to pump and feed^26,43^. The assayed larvae therefore probably preceded these developmental milestones. Their response shows that acoustic exposure can affect behaviour before these specialised structures are expected to be functionally mature, but it does not identify the sensory pathway involved. Direct measurements of particle motion, combined with targeted sensory perturbations, will be required to resolve the underlying mechanism.

In contrast, juvenile and adult free-swimming responses were distributed across a broader range of behavioural features. This should not be interpreted as reduced sensitivity; rather, it reflects the increasing complexity of the locomotor repertoire during the life cycle. In juveniles, the two noise treatments produced qualitatively contrasting effects on trajectory organisation. MSFD-NB noise resulted in more directional and less irregular swimming paths while Vessel-DNS noise generated more convoluted trajectories. Because juvenile swimming is more sustained and spatially organised than larval swimming, these opposing responses suggest that different acoustic signals disrupt the balance between directed locomotion, exploration and local reorientation in distinct ways.

Free-swimming adults exhibited the broadest qualitative behavioural changes, though with substantial variability among individuals, meaning that no comparison remained significant after correction. This observed variability may be consistent with the adult stage possessing the most diverse and context-dependent behavioural repertoire across *O. dioica* ontogeny^27^. As behavioural complexity increases, acoustic treatment in adults appears to be distributed across multiple behavioural dimensions (features) rather than concentrated in a single dominant response. Here, the treatments also diverge: MSFD-NB noise increases turning behaviour while reducing anisotropy and tail-beat amplitude and frequency, consistent with decreased directional stability. Vessel-DNS, however, promotes directionally biased movement, while also altering rotational and propulsive dynamics. Such signal-dependent responses are consistent with findings across a wide range of aquatic organisms, where acoustic treatments differing in spectral and temporal structure often elicit distinct behavioural outcomes^44,45^. Rather than a simple ontogenetic gradient of declining sensitivity, the juvenile and adult data indicate that the same acoustic treatment is expressed through progressively broader and more context-dependent behavioural components as the motor repertoire matures.

Notably, these behavioural changes were not confined to the exposure period in the developmental stages. Larvae and juveniles continued to deviate from controls after treatment had ended, whereas free-swimming adults showed no persistent change, indicating that the effect of noise outlasted the stimulus during development. MSFD-NB noise reorganised motor output in a stage-dependent manner. Larvae had stronger tailbeats and faster turning whereas juveniles showed the opposite shift in propulsion. By contrast, vessel noise suppressed the locomotion of both developing stages. Larvae swam slower, less directional paths, and juveniles showed weaker, straighter swimming indicating that even a short acoustic exposure can leave a transient behavioural trace once the signal has ceased, and that this trace is itself stage-and signal-specific.

The pronounced larval behavioural sensitivity is consistent with transcriptional responses reported during development^35^. Although those transcriptomic data were obtained from embryos rather than swimming larvae, they support the broader possibility that developmental responses to noise involve neural and neuromuscular pathways. Direct links between embryonic transcriptional changes and the larval movement phenotypes observed here remain to be tested.

### House-dynamics changes in the context of noise exposure

Following our analysis of free-swimming behaviour, we examined whether acoustic exposure also affected adult behaviour inside the house and the filtering function it supports. Movement inside the house is constrained by the mechanical interaction between the animal and its feeding structure, which is essential for particle capture, feeding efficiency and protection. Responses within the house to noise differed from free swimming. Under MSFD-NB noise, adults inside the house displayed increased body curvature and tail-beat amplitude, opposite to the reduced tail activity seen in free-swimming adults. Vessel-DNS noise, in contrast, reduced movement within the house. Interestingly, the timing and number of house expansions were not clearly altered by either treatment, indicating that noise perturbed the animal’s movement within the house without disrupting the construction and renewal of the structure itself.

These house-associated changes could alter filter-feeding performance and clearance rate, although neither was measured here. Our results therefore motivate future laboratory and mesocosm experiments that combine behavioural measurements with direct assays of feeding, clearance and house production to determine whether noise exposure affects appendicularian contributions to carbon flux.

### Broader ecological and mechanistic context

The changes described above can be placed in a broader ecological context. Since vertical positioning in the water column depends on the same directed, coordinated swimming assessed throughout this study, the juvenile and adult locomotor changes reported here could plausibly extend to diel vertical migration as well, a well-documented behaviour in this species^46^, where the animals migrate to depth during the day, largely to avoid visually hunting predators, and rise back towards the surface at night to feed on the higher phytoplankton concentrations found there. This could have primary consequences for vertical carbon flux and pelagic food-web structure, though these links remain to be tested directly.

Moreover, a recent study of predator-evoked behaviour in the same species^47^ showed that adult *O. dioica* exposed to a copepod predator showed a similar state-level reorganisation as seen under noise, and that this response required the physical presence of the predator, not chemical cues alone. The sensitivity of adults to mechanically generated predator cues make particle motion a plausible candidate stimulus, but the present experiments do not distinguish particle motion from sound pressure or substrate-transmitted vibration.

However, predator exposure delayed house expansion and reduced turnover, whereas noise left expansion unchanged and altered in-house pumping instead, indicating that noise acts on the fine motor control of filtration rather than on the decision to rebuild a house. Nevertheless, under natural conditions, anthropogenic noise may interfere with behavioural processes that are also critical for feeding and predator avoidance.

Our findings show that playback treatments alter several components of the stereotyped behavioural repertoire of *O. dioica*, with effects differing among life stages, behavioural contexts and exposure signals. Adults also changed their movement within the feeding house, although house-expansion latency was not detectably affected. These observations establish behavioural sensitivity across the life cycle but do not by themselves demonstrate changes in feeding performance or carbon export. Future experiments combining calibrated pressure and particle-motion measurements with simultaneous behavioural, clearance and house-production assays will be required to quantify the functional and ecological consequences.

## STUDY LIMITATIONS

Several methodological limitations should be considered when interpreting these findings. First, each acoustic treatment was delivered at a single intensity and with a fixed temporal structure. Consequently, we cannot determine how the observed stage-and signal-specific responses vary with exposure to intensity, duration or intermittency. Establishing such dose– response relationships will be essential for informing ecologically relevant risk assessments and regulatory thresholds.

Second, our experimental assays necessarily simplified the natural environment experienced by *O. dioica*. The restricted recording depth prevented us from studying depth-dependent behaviours such as diel vertical migration, which we identify as a plausible ecological consequence of the locomotor changes reported here but did not test directly. In addition, although comparisons with predator-response data suggest a potentially shared mechanosensory basis, noise exposure and predation were not tested simultaneously, leaving potential interactions between these stressors unresolved.

Finally, our two noise treatments differ from one another in intensity and spectral content, and additionally in the separate exposure settings, so the present design cannot attribute the diverging behavioural outcomes to any single acoustic parameter. Disentangling which specific properties of each signal drive these effects would require a broader set of controlled acoustic manipulations. In any case, the results demonstrate that *O. dioica* behaviour is not only affected by high-intensity playbacks, but also by vessel noise at lower intensities, addressing one of the aims of this work.

## RESOURCE AVAILABILITY

### Lead contact

Further information and requests for resources and reagents should be directed to and will be fulfilled by the lead contact, M. Chatzigeorgiou

### Materials availability

This study did not generate any new materials.

### Data and code availability

Raw video data will be deposited in Zenodo upon publication.

## ACKNOWLEDGMENTS

This research was funded by JPI Oceans, Project NoiseInTheSea-2022-0011 “Characterization of maritime noise in different european basins and its impact on ecological relevant deuterostome invertebrates” by L.M., G.S., S.C., G.B, R.P., E.S., M.C., F.R., A.S., A.T. C.C., R.M.A.P., I.G, G.Z. We further acknowledge funding from the Michael Sars Centre core budget (NFR grant 234817 “Sars International Centre for Marine Molecular Biology Research, 2013–2022”), grants from the Research Council of Norway to M.C. (339399 and 335582). This work was supported by a Research Grant from HFSP (Ref.-No: RGP007/2026) to M.C. with the award DOI https://doi.org/10.52044/HFSP.RGP0072026.pc.gr.243781. CC and ERQ were funded by PCI2022-135017-2, PID2022-141627NB-I00, PID2025-170547NB-I00 and RED2024-153825-T from Spanish Ministerio de Ciencia, Innovación y Universidades MICIU/AEI/10.13039/501100011033 and by ERDF/EU. ERQ also received an EMBO Scientific Exchange Grant for a research visit to the Michael Sars Centre. CC was supported by ICREA Acadèmia Ac2215698 and 2025 SGR 00782 from AGAUR, Generalitat de Catalunya. R.M.A.P and I.N.P. thank AGAUR (generalitat de Catalunya) for the funding of HER (Human-Environment Research) with code 2021 SGR01396. R.P. S.C. and L.M were funded by MUR, Project PRIN 2022 Prot. 2022SF7HY9 “NoDe: The impact of anthropogenic noise on marine invertebrate deuterostomes: development, behavior and resilience”.

## AUTHOR CONTRIBUTIONS

Conceptualization: E.R.Q., C.C. and M.C.; methodology: E.R.Q., O.T., S.N., R.d.S.M.B. and M.C.; data curation: E.R.Q., O.T., R.d.S.M.B.; investigation: E.R.Q., O.T.; visualization: E.R.Q., O.T. and M.C.; resources: C.C., A.E.A., D.M.C., M.C.; funding acquisition: C.C., D.M.C. and M.C.; project administration: C.C. and M.C.; supervision: C.C. and M.C.; writing – original draft: E.R.Q., O.T. and M.C.; writing – review and editing: E.R.Q., C.C., O.T., S.N., R.d.S.M.B., V.Z.-C., I.N.-P., L.M., G.S., S.C., M.B., E.S., R.P., G.B., A.E.A., D.M.C., G.Z. and M.C.

## DECLARATION OF INTERESTS

The authors declare no competing interests.

## EXPERIMENTAL MODEL AND STUDY PARTICIPANT DETAILS

Wild *O. dioica* were collected in coastal waters near Bergen, Norway, and cultivated for several generations at the Michael Sars Centre, University of Bergen, Norway, before use. The animals were kept according to standard protocol at 12 ± 0.5 °C resulting in a 7-day life cycle [24].

## METHOD DETAILS

### Noise exposure and set-up experimental calibration

A glass tank aquarium measuring 60 cm long, 30 cm wide and 30cm high was filled with 29 L of water, and an AKUASOUND_16 underwater speaker (MAUSOUND, San Damaso, Modena) connected to a custom-made amplifier (Akuasound, Mausound) equipped with a USB input for playback of the acoustic signals stored on a USB flash drive. *O. dioica* animals were placed in a 2 L glass beaker filled with 1.8 L of seawater suspended from a nylon rod attached to the ceiling of the tank in the middle of the water column to avoid the transmission of vibration from the walls of the tank to the beaker (**Fig. 2A**). The acoustic environment of the tank was characterised to ensure accurate and reproducible noise conditions by measuring sound pressure levels using an AS-1 hydrophone (Aquarian Audio, Anacortes, WA, USA) connected via BNC interface to a data acquisition module (SpectraDAQ-200), interfaced through USB 3.0 to a computer running SpectraPLUS-SC software. Signals were digitized at a sampling rate of 44.1 kHz with 16-bit resolution. The tanks and the playback signals used in the study (MSFD-NB and Vessel-DNS) were acoustically characterized and corrected following the methodology described in Zaffaroni-Caorsi et al^31^. Exposure occurred at a selected standardized point for all the post-exposure experiments (Table S1). All reported acoustic levels are expressed in dB re 1 μPa.

Experiments recorded during noise exposure assays were performed directly within the behavioural arena, and the set-up (Fig. 1A) was characterised as well. Treatment was delivered through an airborne loudspeaker Control 1 Pro Two-Way Professional Compact Loudspeaker System positioned outside the water-containing behavioural chamber and connected to an AMP 2 DG amplifier (Auna) and a computer-based playback system. Playback levels were measured using a Class 1 sound level meter (Larson-Davis LD831-C) with Z-frequency weighting, with the microphone positioned approximately 1 cm above the area of the behavioural arena occupied by the animals, as close as possible to their location, to standardise stimulus delivery across experiments. Because acoustic pressure was not measured directly within the water of the behavioural arena, airborne measurements cannot be interpreted as the underwater sound-pressure levels experienced by the animals. For consistency in the presentation of acoustic measurements, airborne sound-pressure levels were additionally expressed using the conventional underwater reference pressure of 1 µPa (**Table S2**).

Airborne and underwater sound-pressure levels use different reference pressures, namely 20 µPa in air and 1 µPa in water, corresponding to a 26 dB difference in the reported sound-pressure level^48^. Accordingly, reference-adjusted values were calculated as:

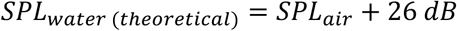

These reference pressure conversion values should not be interpreted as theoretical or direct measurements of underwater sound pressure within the behavioural arena. The actual acoustic field may have been influenced by air–water transmission, chamber geometry, shallow water depth, container walls, reflections, and mechanically transmitted vibrations. This limitation should be considered when comparing these experiments with assays conducted in the standard underwater exposure tank.

### *Oikopleura dioica* behavioural recording

To track *O*. *dioica* at different stages, we placed the animals in ‘agarose arenas’ and used an array of three behavioural setups which we have previously described in^28,29^. Agarose arenas were created using custom 3D-printed PLA moulds^29^. Briefly, 35-mm Petri dishes were filled with ∼9 ml of 0.8% agarose prepared in filtered sea water (FSW). While the agarose was still warm, the moulds were inserted into the agarose and left in place until gelation. Moulds were then removed, excess agarose was trimmed, and the arenas were hydrated with FSW. The resulting circular arenas measured 10 mm in diameter and 3 mm in depth, corresponding to an approximate volume of 240 mm³. The agarose coating of the arena is essential to prevent *O*. *dioica* from sticking to the plastic Petri dish. The animals were recorded at 30 frames/second.

### Recordings during noise exposure

For the larval and juvenile stages, controlled fertilisations were performed as previously described in Marti-Solans et al.^33^. Eggs were fertilized with a sperm pool obtained from a minimum of 3 to 4 males. The first cell division around 20-30 min pf was checked to confirm fertilization and the embryos were kept in separate temperature-controlled refrigerators: larvae at 14 °C and juveniles at 12 °C. These stage-specific temperatures were used to obtain animals at the target developmental ages for each assay until recording.

For the larval stage, fertilized eggs were incubated until 7.5 h pf, then split into two groups and immediately after recorded for a total of 45 min consisting of a 15-minute period without playback, followed by 15 min of noise and then 15 more min without noise.

For the juvenile stage, animals were incubated until 18 h pf at 12 °C and then recorded for a total of 45 min consisting of a 15-minute period without playback, followed by 15 min of noise and then 15 more min without playback.

For the adult stage, day 4 and 5 animals out of a 7-day cycle at 12 °C were selected (**Fig. 1B**). Adults inside their house were selected, forced out of the house and then recorded for a total of 45 min. The animals were exposed to noise during the first 15 min of the recording, then the noise was stopped and the animals were recorded for 30 more min.

### Recordings after noise exposure

For the larval and juvenile stages, controlled fertilisations were performed as previously described above.

For the larval post-exposure assay, fertilised eggs were incubated until 5.5 h post-fertilisation and divided into control and treatment groups immediately after hatching. Groups were transferred to the noise-exposure tanks^31,35^ and exposed for 2 h to either the noise treatment or background noise. Larvae were then recorded for 15 min.

For the juvenile stage, animals were incubated until 18 h pf at 12 °C, exposed to noise^31^ for one hour and then recorded for a total of 45 min.

For the adult stage, day 4 and 5 animals out of a 7-day cycle at 12 °C were exposed to noise as described in^31^ for one hour. Adults inside the house were selected, forced out of the house and then recorded for a total of 15 min.

### Tracking and data processing

Body-part tracking was performed using three stage-specific DeepLabCut (DLC) 3.0.0rc6^49^ models (larva, juvenile, adult) developed in the Chatzigeorgiou laboratory^27^ and applied to the recordings obtained in this study; the models are available at Zenodo (<u>10.5281/zenodo.17473173</u>, 10.5281/zenodo.17459389, 10.5281/zenodo.17491493). Resulting x-y coordinates and likelihood scores for each tracked body part were exported as CSV files for downstream analyses.

Tracking confidence was assessed from DLC likelihood values, and detections with likelihood <0.80 were discarded. Remaining coordinates were filtered to remove extreme outliers using an interquartile-range criterion (IQR threshold = 2.0) and biologically implausible frame-to-frame displacements (>100 pixels). Short isolated tracking segments were excluded by requiring continuous tracking for ≥3 s (90 frames at 30 frames s−1) for larvae and juveniles and ≥1 s for adults. Internal gaps of up to three frames, when bounded by valid detections, were filled by linear interpolation without extrapolation beyond the first and last valid detections. Recordings were acquired at 30 frames s−1 and, unless otherwise stated, trajectories were downsampled fivefold to 6 Hz for downstream behavioural analyses. Tail-beat analysis and trajectory plots were performed at the native frame rate.

Experimental metadata, including developmental stage, treatment, recording setup, animal identity and recording segment, were automatically extracted from standardised filenames and validated for consistency.

### Behavioural metrics extraction

Kinematic features were derived from centre-of-mass (COM) trajectories calculated using trunk body landmarks as described previously^27^. Speed was computed from frame-to-frame COM displacement and converted to µm/s using a calibrated spatial scale (11.56 µm/pixel). Acceleration was calculated as the absolute change in speed per unit time. Heading was defined as the direction of COM displacement, and angular velocity was calculated from frame-to-frame changes in heading, wrapped to (−180°, 180°) and divided by the frame interval. Angular acceleration was derived from changes in angular velocity..

For skeleton analysis, body centrelines were interpolated through trunk and tail anchor points. Skeleton-based descriptors included body curvature, tangent angle, body axis orientation, quirkiness, tail-beat frequency, and tail-beat amplitude. Body curvature and tangent angles were estimated using Savitzky–Golay smoothing, while tail-beat frequency was determined from distal tail displacement using fast Fourier transform (FFT)-based spectral analysis, as described previously^27^.

For skeleton analysis, body centrelines were extracted by interpolating through trunk and tail anchor points. Skeleton-based behavioural descriptors were subsequently derived from the tracked body configuration, including body curvature, tangent angle, body axis orientation, quirkiness, tail-beat frequency, tail-beat amplitude and angular velocity. Body curvature and tangent-angle estimates were obtained from Savitzky-Golay smoothed body outlines, whereas tail-beat frequency was estimated from the lateral displacement of the distal tail using fast Fourier transform (FFT)-based spectral analysis as described in Tolstenkov et al^27^.

Behavioural states were classified as quiescence, dwelling or roaming based on centre-of-mass (COM) speed. Speed traces were resampled to 6 Hz and smoothed using a 10-frame running mean. Activity bouts were detected using stage-specific peak and base speed thresholds of 0.10 and 0.07 mm/s for larvae and juveniles, and 0.60 and 0.40 mm/s for adults, respectively. Bout boundaries were determined from crossings of the quiescence baseline, with a maximum duration of 30 s; bouts separated by ≤0.5 s were merged. Activity bouts were classified as dwelling or roaming based on whether their peak speed was below or at least equal to the mean speed of stage-matched control recordings. Remaining valid frames were classified as quiescent. State occupancy, bout frequency and bout-duration statistics were calculated for each individual.Arena and path-shape metrics were derived from COM trajectories using sliding 3-s windows. Net displacement was defined as the Euclidean distance between the first and last positions, and tortuosity as the ratio of total path length to net displacement. Radius of gyration was calculated as the root mean square distance from the window centroid, while ringness was defined as the coefficient of variation of radial distances. Anisotropy was calculated as the ratio of the larger to the smaller eigenvalue of the coordinate covariance matrix, and path entropy as the Shannon entropy of its normalised eigenvalues. Winding was calculated from the absolute change in unwrapped movement direction divided by 2π. Tortuosity was retained only for windows with net displacement ≥5 pixels and non-zero path length. Moment-based metrics required at least four valid positions, whereas net displacement, tortuosity, winding and path entropy required complete windows. House feature: The cellulose house is essential for *Oikopleura* feeding and survival. Thus, after the video recordings of the adults, the stereotypical movement implicated in the new house expansion was manually annotated for the first 15 min of recording, for the different noise exposures. The time was registered in the form of latency (seconds) until house expansion, with a cutoff of 900 seconds.

### Dimensionality reduction

Candidate features comprised per-animal summaries of kinematic, trajectory-shape, skeleton-derived and behavioural-state metrics. Feature filtering was performed independently within each developmental stage. Features missing in more than 30% of animals were excluded, followed by removal of near-constant features with a coefficient of variation below 0.20. Redundancy was reduced using absolute Spearman correlations, with thresholds of 0.75 within feature categories and 0.92 across categories. To preserve representation of each behavioural domain, minimum quotas of three kinematic, two arena, two skeleton and two behavioural-state variables were applied.

Principal component analysis (PCA) was restricted to recordings obtained during noise exposure and performed separately for larvae, juveniles and adults recorded inside and outside the house. Within each stage, MSFD-NB and Vessel-DNS datasets, including their respective control and noise groups, were analysed together. PCA was implemented in scikit-learn, retaining up to six components based on scree plots and cumulative explained variance, targeting 85%. Component scores, loadings and explained variance were used for visualisation, with 95% confidence ellipses included in score plots. Explained variance and component loadings are presented in Fig. S1. Retained features were used for percentage-change summaries (Figs. 1 and 2B) and multivariate comparisons.

### Quantification and statistical analysis

Following quality control, behavioural analyses included 1198 animals: 429 larvae, 526 juveniles and 243 adults. Of these, 150 larvae, 208 juveniles and 120 adults were analysed in the during-exposure assay, and 279 larvae, 318 juveniles and 123 adults in the after-exposure assay. The final dataset comprised approximately 62.5 million tracked frames (578 h at 30 frames s⁻¹), distributed across larvae (173.6 h), juveniles (227.8 h) and adults (177.0 h). Treatment-and stage-specific sample sizes are reported in Supplementary Table S2.

Behavioural metrics were summarised per animal across valid recording segments using the mean, median and 90th percentile. Per-animal means were used for feature selection, principal component analysis (PCA) and percentage-change comparisons (Figs. 1 and 2B), whereas the 90th percentile was used for post-exposure comparisons (Fig. 3).

Each noise-exposed group was compared with its corresponding control within the same life stage, assay, and noise type using two-sided Mann–Whitney U tests. Effect sizes were reported as percentage changes in group means, Cohen’s d and Cliff’s δ, with 95% bootstrap confidence intervals for Cliff’s δ (2,000 resamples). P-values were adjusted using the Benjamini–Hochberg (BH) procedure across metrics within each stage × assay × noise type combination, including PCA-retained metrics (Figs. 1 and 2B), all 19 metrics (Fig. 3) and metrics included in the Cliff’s δ overview (Fig. S3).

Multivariate differences were quantified as Euclidean distances between noise and control centroids in z-scored, PCA-retained feature space, with missing values imputed using pooled medians independently of treatment labels. Significance was assessed using 10,000 label permutations, with bootstrap ranges from 2,000 resamples. The analysis was repeated for PC1–PC2, PC3–PC4 and Kaiser-retained components. P-values were BH-adjusted across groups separately for each noise type, representing independent experiments. Centroids with 95% bootstrap confidence ellipses are shown in Fig. S2.

Box plots (Fig. 3) show the median and interquartile range, with whiskers at the 5th and 95th percentiles. Test statistics, effect sizes and unadjusted and BH-adjusted p-values are provided in Supplementary Tables S2–S13. Statistical significance was assessed at α = 0.05, using adjusted p-values where applicable.

### Software

All analyses were performed using a custom analysis pipeline written in Python 3.11.14, using NumPy 2.3.5, pandas 2.3.3, SciPy 1.16.3, scikit-learn 1.7.2, statsmodels 0.14.5, joblib 1.5.2, openpyxl 3.1.5, and Matplotlib 3.10.8. The analysis pipeline is available at https://github.com/ChatzigeorgiouGroup/Oikopleura_noise_behavior.

## Notes

### Competing Interest Statement

The authors have declared no competing interest.

