## Supplementary Figures 1-3 and Tables S1 and S2 for "Anthropogenic noise affects *Oikopleura dioica* behaviour in a life stage- and signal-dependent manner"

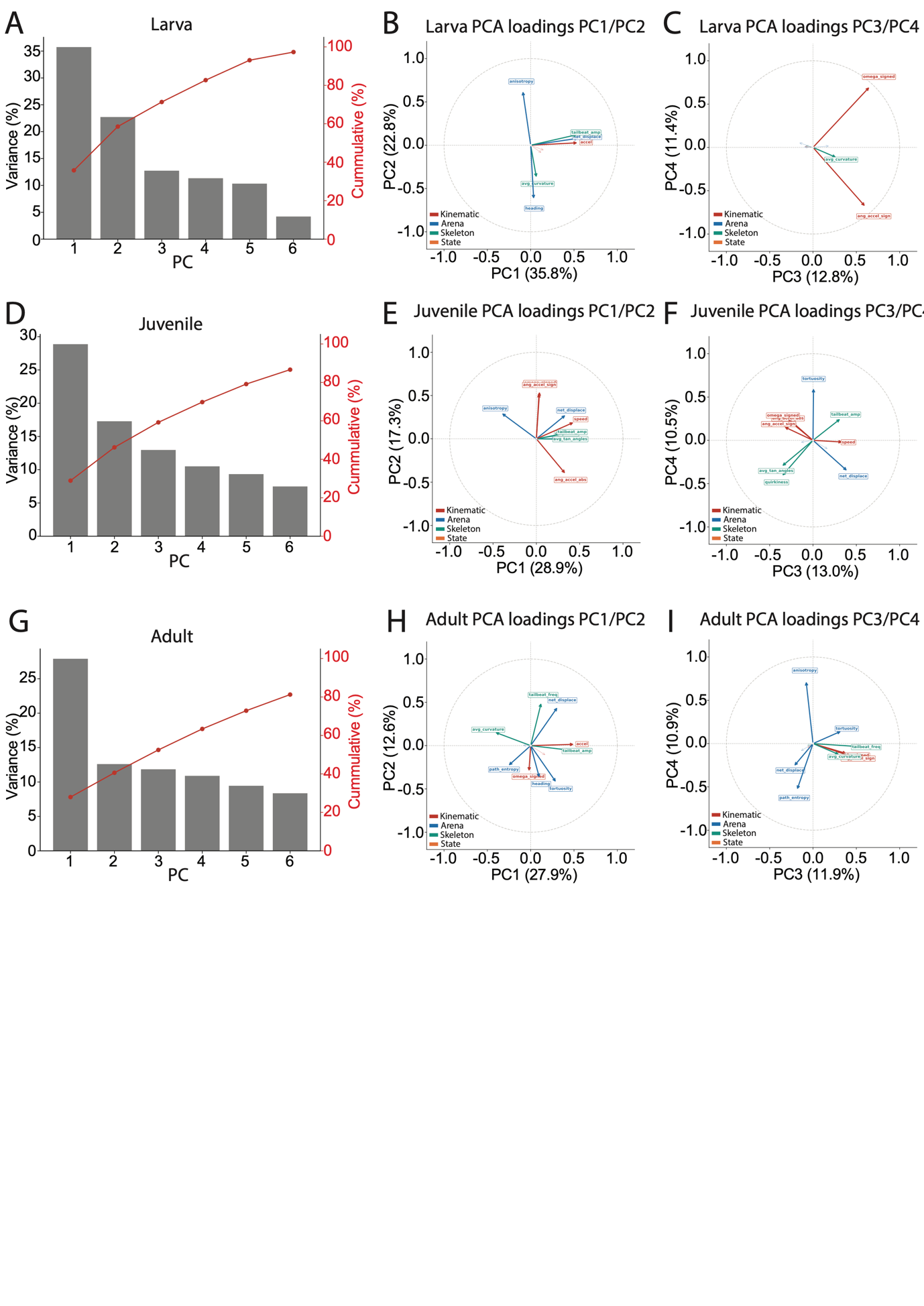

**Figure S1**. **Principal component analysis of the behavioural metrics, by life stage (larva, juvenile, adult). (A, D, G)** Scree plots showing the variance explained by each principal component (bars) and the cumulative variance explained (line) for larvae, juveniles and adults, respectively. **(B, C, E, F, H, I)** PCA loading plots showing how each behavioural metric contributes to PC1–PC4, coloured by parameter category (kinematic, arena, skeleton, and behavioural state). Please also see Tables S10 and S12.

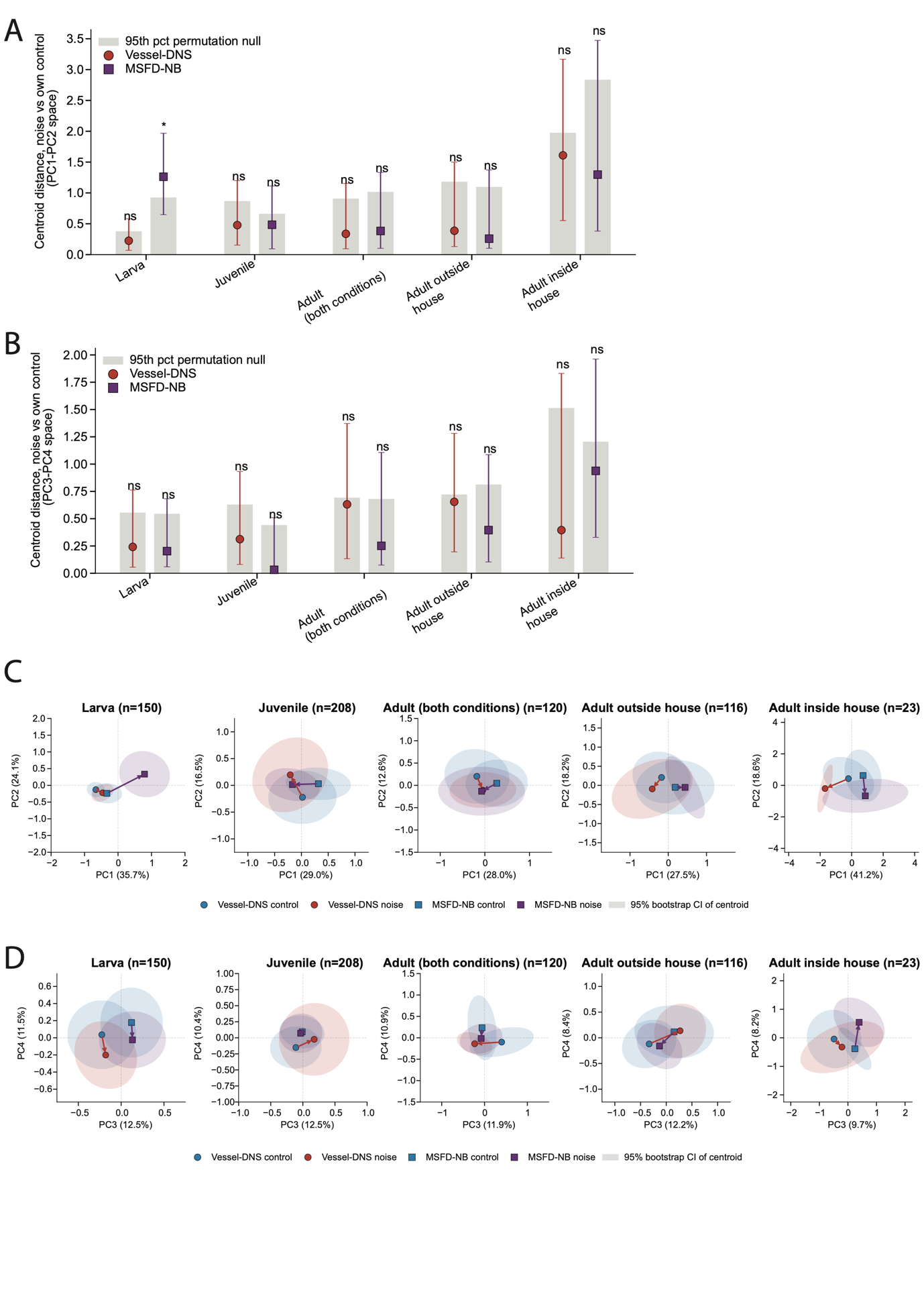

**Figure S2**. Multivariate behavioural responses to MSFD-NB and Vessel-DNS exposure across developmental stages and house conditions. (A, B) Euclidean distances between noise-exposed and control group centroids in the PC1–PC2 and PC3–PC4 spaces, respectively, for larvae, juveniles and adults (all adults, outside and inside the house). Red circles indicate Vessel-DNS and purple squares MSFD-NB treatment. Error bars show 95% bootstrap ranges (2,000 resamples), and grey bars indicate the 95th percentile of the permutation null distribution (10,000 permutations). Asterisks indicate significant differences following Benjamini–Hochberg (BH) correction (adjusted p < 0.05); ns, not significant. (C, D) PCA projections in PC1–PC2 and PC3–PC4 spaces, respectively. Symbols indicate group centroids, coloured regions show 95% bootstrap confidence ellipses, and lines connect noise-exposed and corresponding control centroids. Sample sizes are indicated above each plot. Percentages on the axes indicate variance explained by each principal component. Please also see Table S12.

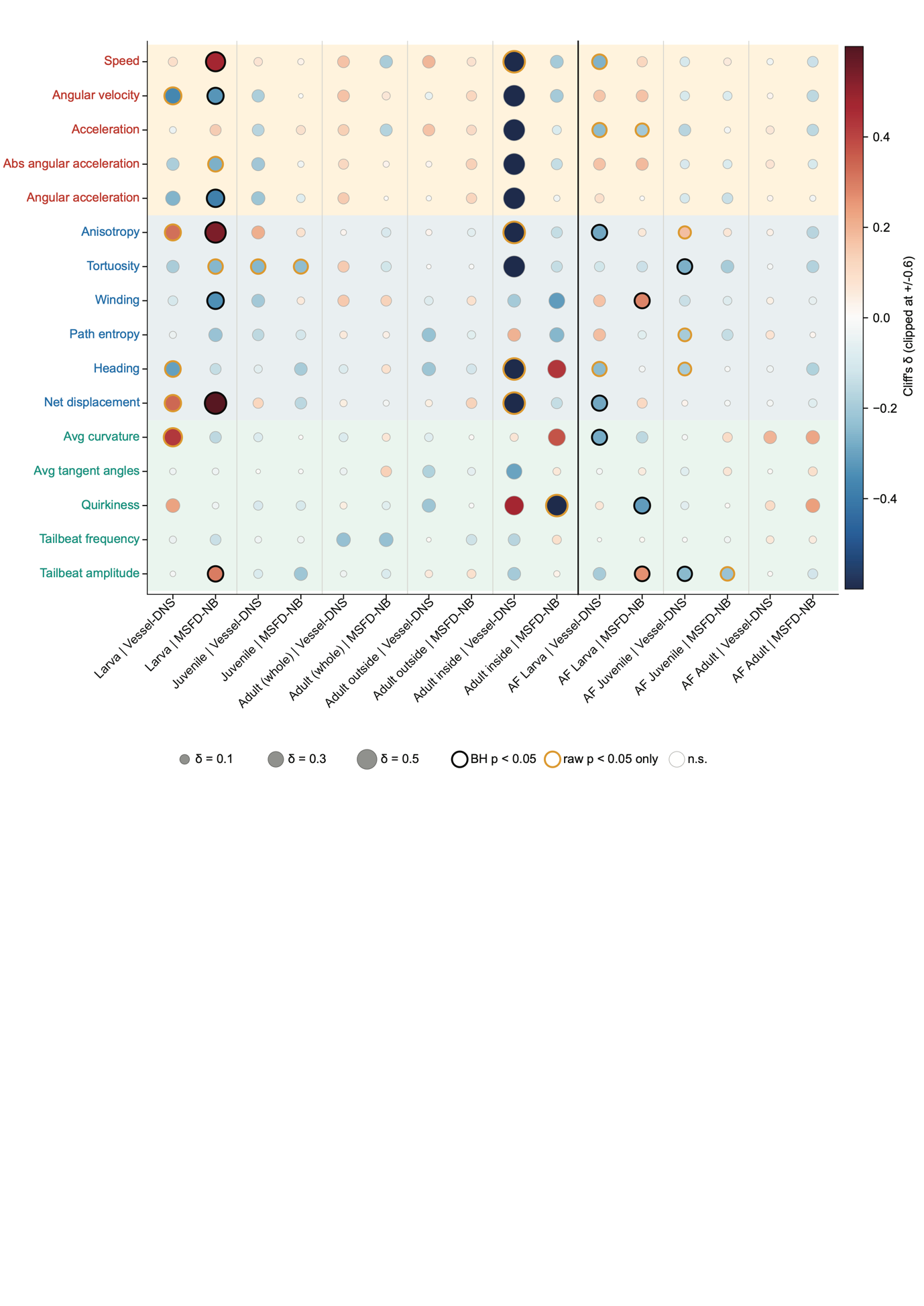

**Figure S3**. Effect sizes and statistical significance of behavioural responses to MSFD-NB and Vessel-DNS exposure across developmental stages. Cliff’s δ effect sizes for kinematic, arena, and skeleton metrics in larvae, juveniles, and adults (including adults inside and outside the house) during exposure, and in larvae, juveniles, and adults after exposure (AF). Circle colour indicates the direction and magnitude of the effect (red, higher values under noise; blue, lower values), and circle size represents the absolute effect size. Black outlines indicate significant differences after Benjamini–Hochberg (BH) correction (adjusted p < 0.05), orange outlines indicate nominal significance only (uncorrected p < 0.05), and unoutlined circles indicate non-significant differences. Behavioural metrics are grouped by category (kinematic, arena and skeleton). Statistical comparisons were performed using two-sided Mann–Whitney U tests, with BH correction applied across metrics within each developmental stage, assay, and noise treatment. Sample sizes and statistical results are provided in Table S13

**Table S1**. Standardised treatments for post-exposure experiments. *SPL, Sound Pressure Levels.*

| **Treatment** | **Spectral Composition** | **Signal Type** | **SPL** *(dB re 1 µPa)* |
| --- | --- | --- | --- |
| **Control** | Background noise | Ambient background | 83 |
| **MSFD-NB** | 63-125 Hz band with dominant energy peaks at 63 and 125 Hz | Standardised | 158 |
| **Vessel-DNS** | 100-1000 Hz band with dominant energy peaks at 63 and 125 Hz | Vessel recording | 130 |

**Table 2. Airborne playback levels measured close to the behavioural arena and corresponding theoretical underwater reference levels.** Maximum unweighted equivalent sound-pressure levels (LZeq) were measured in air close to the behavioural arena during the level-60 playback setting. Reference-adjusted equivalent values were obtained by expressing airborne levels relative to the conventional underwater reference pressure of 1 µPa.

| **Acoustic treatment** | **Measured airborne playback level, LZeq** | **Reference-adjusted equivalent value** |
| --- | --- | --- |
| **Ambient room level** | 79.1 dB re 20 µPa | 105.1 dB re 1 µPa |
| **MSFD-NB** | 102.8 dB re 20 µPa | 128.8 dB re 1 µPa |
| **Vessel-DNS** | 94.0 dB re 20 µPa | 120.0 dB re 1 µPa |
